# Variation in *Leishmania* chemokine suppression driven by diversification of the GP63 virulence factor

**DOI:** 10.1101/2021.02.01.429140

**Authors:** Alejandro L. Antonia, Amelia T. Martin, Liuyang Wang, Dennis C. Ko

**Author notes:** To whom correspondence should be addressed: Dennis C. Ko, 0049 CARL Building Box 3053, 213 Research Drive, Durham, NC 27710. @denniskoHiHOST.

## Abstract

Leishmaniasis is a neglected tropical disease with diverse infection outcomes ranging from self-healing lesions, to progressive non-healing lesion, to metastatic spread and destruction of mucous membranes. Although resolution of cutaneous leishmaniasis is a classic example of type-1 immunity leading to well controlled self-healing lesions, an excess of type-1 related inflammation can contribute to immunopathology and metastatic spread of disease. *Leishmania* genetic diversity can contribute to variation in polarization and robustness of the immune response through differences in both pathogen sensing by the host and immune evasion by the parasite. In this study, we observed a difference in parasite chemokine suppression between the *Leishmania (L*.) subgenus and the *Viannia (V*.) subgenus, which is associated with severe immune mediated pathology such as mucocutaneous leishmaniasis. While *Leishmania (L*.) subgenus parasites utilize the virulence factor and metalloprotease glycoprotein-63 *(gp63)* to suppress the type-1 associated host chemokine CXCL10, *L. (V*.*) panamensis* did not suppress CXCL10. To understand the molecular basis for the inter-species variation in chemokine suppression, we used *in silico* modeling of the primary amino acid sequence and protein crystal structures to identify a putative CXCL10-binding site on GP63. We found the putative CXCL10 binding site to be located in a region of *gp63* under significant positive selection and that it varies from the *L. major* wild-type sequence in all *gp63* alleles identified in the *L. (V*.*) panamensis* reference genome. We determined that the predicted binding site and adjacent positively selected amino acids are required for CXCL10 suppression by mutating wild-type *L. (L*.*) major gp63* to the *L. (V*.*) panamensis* allele and demonstrating impaired cleavage of CXCL10 but not a non-specific protease substrate. Notably, *Viannia* clinical isolates confirmed that *L. (V*.*) panemensis* primarily encodes non-CXCL10-cleaving *gp63* alleles. In contrast, *L. (V*.*) braziliensis* has an intermediate level of activity, consistent with this species having more equal proportions of both alleles at the CXCL10 binding site, possibly due to balancing selection. Our results demonstrate how parasite genetic diversity can contribute to variation in the host immune response to *Leishmania* spp. infection that may play critical roles in the outcome of infection.

## Introduction

Parasites in the genus *Leishmania* infect over 1.6 million people annually causing a diverse collection of diseases ranging from visceral systemic illness to simple self-resolving cutaneous lesions to diffuse non-healing lesions with metastatic spread(1, 2). This diverse spectrum of disease outcomes is in part mediated by parasite genetic diversity influencing the host-immune response. Due to the high psychological and social impact caused by disfiguring skin lesions(3), and current drug options limited by high prices and severe side effects(4), improved understanding of the mechanisms underlying differential disease outcome is paramount.

Although cutaneous leishmaniasis canonically requires T-helper 1 polarization for lesion resolution, improved understanding of the parasite diversity has led to the elucidation of multiple exceptions to this rule. Experiments using the murine model of leishmaniasis defined the T-helper (T_h_) cell polarization dichotomy where T_h_1 polarization in C57BL6 mice protects against cutaneous leishmaniasis and T_h_2 polarization in BALB/c mice leads to progressive non-healing infections(5, 6). Similarly, humans with self-healing lesions have higher levels of T_h_1 associated cytokines(7-9). However, studies in mice and humans have revealed instances where type 1 immune responses are not protective against cutaneous leishmaniasis. For example, mice infected with the *L. (L*.*) major* Seidman strain develop non-healing lesions despite robust T_h_1 polarization(10, 11). Additionally, patients infected with parasites belonging to the *Viannia* subgenus of parasites have lesions characterized by significantly elevated expression of type-1 associated cytokines such as *IFN*-*γ, Granzyme-B* and *CXCL10(12-14)*. Further, these markers of type-1 associated immunopathology are exacerbated by infection of the parasite with the *Leishmania* RNA virus (LRV1)(15) or co-infection of the mouse with lymphocytic choriomeningitis virus (LCMV) (16, 17). These exceptions to the rule of protective type-1 immunity raise the question: what determines if a type-1 immune response protects the patient or exacerbates disease?

Clues to answering this question may come from understanding *Leishmania* strategies to evade the host immune response(18). *Leishmania* spp. parasites evade the host immune response by diverse mechanisms including inhibition of phagolysosome development, antigen cross-presentation, and intracellular macrophage signaling (reviewed in Gupta et al. 2013(19)). One virulence factor involved in multiple mechanisms of immune evasion is the matrix-metalloprotease glycoprotein-63 (*gp63*)(20) which is able to cleave host substrates involved in anti-parasitic defense including complement C3b(21), the myristolated alanine rich C kinase substrate (MARCKS)(22), and CXCL10(23). These mechanisms interfere with the host response after infection thereby modulating the balance between protection and pathogenesis.

Parasite genetic diversity also drives differences in disease outcome through both host pathogen sensing and pathogen immune evasion. Although there is a high degree of conservation and synteny between parasite strains, differences in genetic structural elements such as gene duplications and losses, transposable elements, and pseudogenes are thought to play a role in mediating distinct disease outcomes(24, 25). These differences are highlighted within the *Viannia* subgenus, which is associated with lesions characterized by host-cytotoxic immunopathology. Differences in the structure of the lipophosphoglycans produced by *L. (V*.*) braziliensis* and *L. (L*.*) infantum* lead to different patterns of toll-like receptor (TLR) activation(26). Beyond sensing, parasite genetic variation facilitates the generation of diverse immune evasion strategies. For example, parasites in the *Viannia* subgenus have undergone a significant expansion in copy number corresponding with an increase in genetic variation of *gp63(27-30)*. This diversification correlates with variation in *gp63* expression and the downstream effect of phagolysosome maturation between different *L. (V*.*) braziliensis* isolates(31)^31^. Furthermore, evolutionary studies have identified a region of positive selection around the active site hypothesized to alter substrate specificity(32, 33). However, specific genetic polymorphisms responsible for driving observed phenotypic differences between parasites that cause distinct forms of disease have yet to be established.

In this study, we characterize how genetic diversity results in variation in function of the glycoprotein-63 virulence factor between the *Leishmania* and *Viannia* subgenera. We observed that *Viannia* parasites have reduced capacity to cleave CXCL10 by GP63, highlighted by complete loss of cleavage by *L*. (*V*.) *panamensis*. To define how interspecies *gp63* variation results in loss of CXCL10 cleavage, we used a combination of modeling protein-protein interactions and identification of sites under evolutionary pressure. First, protein-protein modeling of the GP63-CXCL10 interaction revealed a putative CXCL10 binding site that is mutated from D463 to N463 preferentially in the *Viannia* subgenus. Screening additional *Viannia* species demonstrated that parasites containing both alleles have an intermediate CXCL10 suppression phenotype. Subsequently, using the mixed effects model of evolution (MEME) we found that of 4 out of 79 individual amino acids under episodic positive selection are located within 5 amino acid residues of the predicted CXCL10 binding site. Finally, site-directed mutagenesis of *L*. (*L*.) *major gp63* at the predicted CXCL10 binding site (D463N), as well as adjacent residues under significant episodic positive selection, confirmed this region is involved in cleavage of CXCL10 but not the nonspecific substrate azocasein. Understanding how genetic diversity of GP63 and other virulence factors lead to differences in immune phenotypes between divergent *Leishmania* spp. will be critical in the design and proper application of novel therapies for the prevention and treatment of leishmaniasis.

## Results

### Glycoprotein-63 produced by L. (V.) panamensis (PSC-1) does not cleave human CXCL10

As *Viannia* parasites induce high c*xcl10* expression(12, 13, 15) and have a large copy number expansion(27, 29) of the CXCL10-cleaving protease GP63(23), we asked whether *Leishmania Viannia panamensis* infection results in a net increase or decrease of CXCL10 protein. PMA-differentiated THP-1 macrophages infected with *L*. (*L*.) *major* induced host *cxcl10* transcription but completely suppressed the induction at the protein level (Fig. 1A, B), consistent with our previous observations(23). In contrast, despite the large copy number expansion of *gp63* in *Viannia* parasites, *L*. (*V*.) *panamensis* infection resulted in higher induction of *cxcl10* transcript and no suppression of CXCL10 protein (Fig. 1A, B).

**Figure 1.**
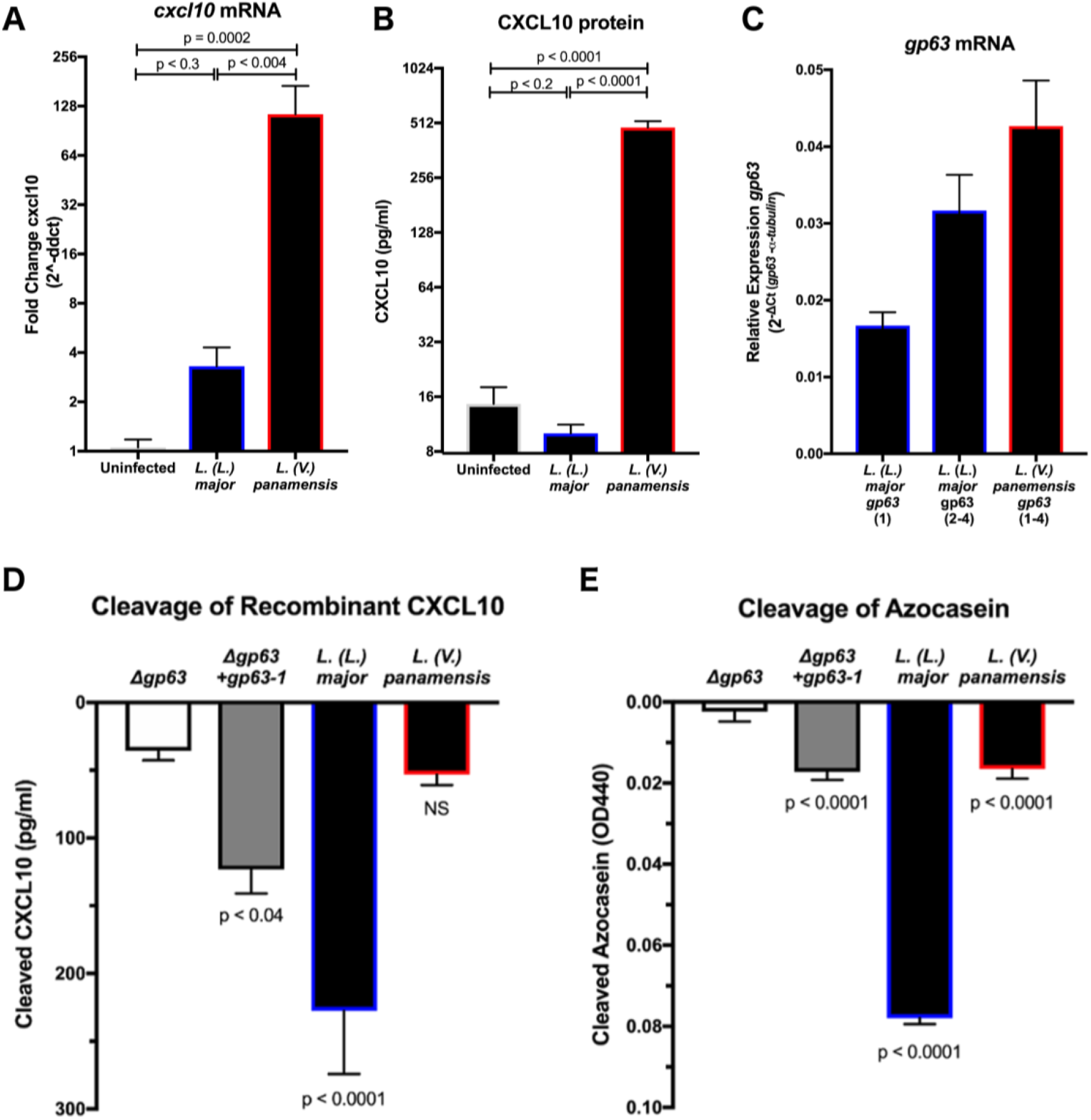
*Leishmania Viannia panamensis* lacks glycoprotein-63 dependent cleavage of human CXCL10. *(A)* L. (L.) major *and* L. (V.) panamensis *infection results in increased CXCL10 transcript in THP-1 monocytes*. PMA differentiated THP-1 monocytes were infected with *L. (L*.*) major* Friedlin or *L. (V*.*) panamensis* PSC-1 at an MOI of 10 for 24 hours. mRNA was quantified by RT-PCR for human *CXCL10* relative to *rRNA45s5* housekeeping gene by ΔΔCt. Average fold change (2^-ΔΔCt^) +/-standard error of the mean is plotted for 8-9 biological replicates across three independent experiments. P-values calculated by one-way ANOVA with Holm-Sidak post-hoc test on C_t_ values. *(B)* L. (L.) major *but not* L. (V.) panamensis *infection suppresses CXCL10 protein in THP1 monocytes*. CXCL10 protein was measured from the supernatants of infected PMA differentiated THP-1 monocytes 24 hours post infection by ELISA. Samples below the ELISA range were set to the lower limit of detection (7.8125 pg/ml). Mean +/-standard error of the mean is plotted for 8-9 biological replicates across three independent experiments. P-values calculated by one-way ANOVA with Holm-Sidak post hoc test on log_2_([CXCL10]). *(C)* L. (L.) major *and* L. (V.) panamensis *both express* gp63 *mRNA at the time of infection*. Parasite mRNA was obtained from parasites incubated at 37°C for 1 hour. mRNA was quanitified by RT-PCR for *Leishmania gp63* relative to α-tubulin housekeeping gene by ΔΔC._t_For primer design to quantify *gp63* expression in different species: *L. (L*.*) major* Friedlin gp63 required two sets of primers to amplify four copies of *gp63* due to sequence variation, whereas *L. (V*.*) panamensis* required a single set of primers due to greater homology between four unique copies. Relative expression (2^-ΔΔCt^) is plotted as mean +/-standard error of the mean for three biological replicates across three independent experiments. *(D)* L. (L.) major *but not* L. (V.) panamensis *cleaves human recombinant CXCL10*. 1×10^6^ parasites were incubated with 500pg/ml of human recombinant CXCL10 at 37°C for 1 hour prior to measuring the remaining CXCL10 by ELISA. Cleaved CXCL10 was calculated by subtracting the CXCL10 in the parasite conditions from a no-parasite media control. Mean cleaved CXCL10 +/-standard error of the man is plotted from 10 biological replicates across 5 independent experiments. *(E)* L. (L.) major *and* L. (V.) panamensis *both cleave the non-specific colorimetric protease substrate azocasein*. 5×10^7^ parasites were incubated with 50mg/ml of azocasein at 37°C for 5 hours. Cleaved azocasein was determined by subtracting OD440 from the no-parasite media control. Mean OD440 of cleaved azocasin +/-standard error the mean is plotted from 7 biological replicates across 5 independent experiments. For (D) and (E), P-values calculated by one-way ANOVA comparing all samples to the *gp63* negative control by Holm-Sidak post-hoc test.

An inability of *L. (V*.*) panamensis* to overcome the increased *cxcl10* transcript could be due to either a lack of *gp63* expression or a reduction in GP63 cleavage activity. To compare *gp63* expression, we measured *gp63* mRNA by *L*. (*V*.) panamensis and *L. (L*.*) major* in cultured promastigotes at the time of infection. The two *Leishmania* spp. expressed comparable *gp63* mRNA (Fig. 1C), suggesting that differences in gene expression do not account for the observed lack of CXCL10 suppression by *L. (V*.*) panamensis*. To compare the ability of *L*. (*V*.) *panamensis* and *L. (L*.*) major* to cleave equivalent amounts of recombinant CXCL10 and control for variation in host chemokine production, we incubated parasites with human recombinant CXCL10. *L. (L*.*) major* that is *gp63* deficient (Δ*gp63*) and complemented (Δ*gp63+gp63-1*) were included as controls. Wildtype *L*. (*L*.) *major* cleaved the majority of CXCL10 by 1 hour; however, there was no significant reduction of CXCL10 by *L*. (*V*.) *panamensis* relative to the Δ*gp63* strain (Fig. 1D). Next, we tested whether this was due to a complete loss of proteolytic activity or a change in substrate-specific activity by repeating the cleavage assay with the non-specific colorimetric substrate azocasein. *L*. (*V*.) *panamensis* cleaved significantly more azocasein than the Δ*gp63* strain, although still had reduced activity relative to wild-type *L*. (*L*.) *major* (Fig. 1E). Therefore, the lack of CXCL10 suppression by *L. (V*.*) panamensis* is due to loss of substrate-specific enzymatic activity of GP63.

### Identification of predicted CXCL10 binding site on GP63

We hypothesized that *gp63* sequence diversity drives the phenotype of reduced CXCL10 cleavage by *L. (V*.*) panamensis*. To test this hypothesis, we modeled where CXCL10 binds to GP63 relative to known functional residues on the protease. The machine learning approach in xgBoost Interface Prediction of Specific-Partner Interactions (BIPSPI)(34) identified a region of three consecutive amino acids on GP63 (Y461, S462, and D463) in the C-terminal domain distant from the predicted matrix metalloprotease substrate binding pocket(35) (Fig. 2A; complete results summarized in Table S1). The site is also distal from known glycosylation sites involved in protein folding and stability(36), the protease active site(37), secretion signal(38), and putative cysteine switch regulatory element(37) (Fig. 2A). Further, this site varied between *Leishmania* and *Viannia* subgenera based on 54 available full length *gp63* sequences (Fig. 2B; 34 *Leishmania*, 18 *Viannia*, 2 *Sauroleishmania*). Thus, the region identified by BIPSI is distinct from known functional domains of GP63 and is divergent between *Leishmania* and *Viannia* subgenera.

**Figure 2.**
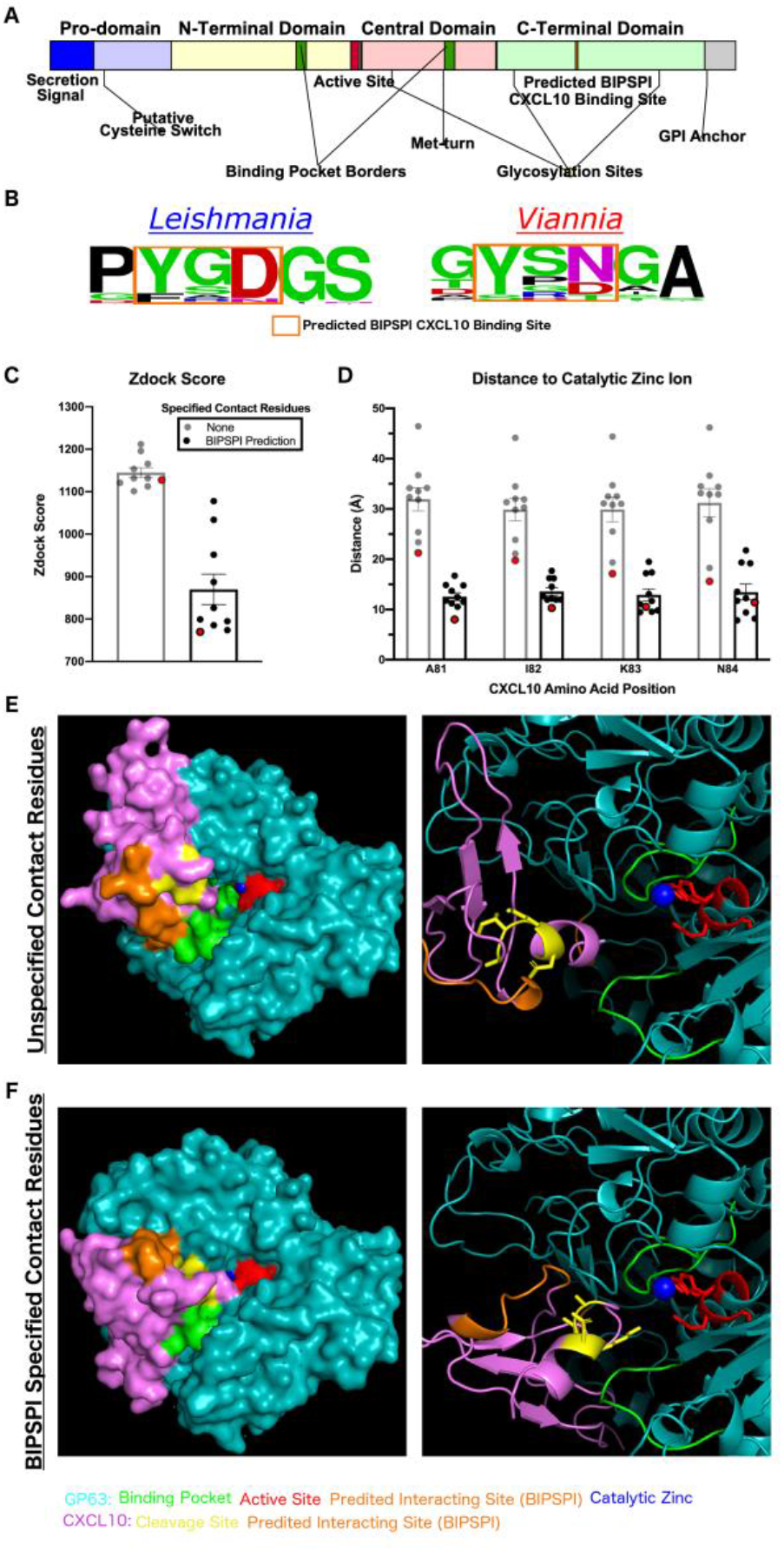
Protein-protein interaction modeling demonstrates a putative CXCL10 binding site on GP63 that closely approximates the GP63 active site and CXCL10 cleavage site. *(A) The predicted CXCL10 binding site is distal from previously described functional residues on GP63*. Domain diagram of GP63 protein summarizing known functional and structural features. *(B) The predicted CXCL10 binding site is nearly invariant among the Leishmania subgenus at position D463 and has been mutated to N463 preferentially in the Viannia subgenera*. 54 homologues of *gp63* from *L. major* (*LmjF_10*.*0460*) were identified by BlastP on TriTrypDB. The sequences represent the *Leishmania* (34), *Viannia* (18), *and Sauroleishmania* (2) subgenera. Multisequence alignment created using ClustalOmega. Sequence logo is shown from AA position 460-465 on the *L. major 10*.*0460* sequence. (C-F) *Modeling of the GP63-CXCL10 interaction with specification of the predicted binding site localizes CXCL10 to the active site and decreases distance to catalytic zinc-ion*. GP63 (PDB entry 1lml) and CXCL10 (PDB entry 1o7y) protein-protein interaction was modeled using Zdock with either no specified contact residues or the BIPSPI predicted binding site specified as contact residues. (C) The mean Zdock score, an energy based scoring function, is plotted for the top 10 model predictions for each condition. (D) The distance from the known cleavage site on CXCL10, in between A81 and I82, to the catalytic zinc ion was measured using PyMol for the top 10 predicted models. Mean distance in angstroms with SEM is plotted. For (C) and (D) the model with the shortest distance between cleavage site and active site is highlighted in red and the model crystal structures shown in (E) and (F). GP63 is shown in in teal and CXCL10 in purple along with annotation of functional residues as follows: GP63 active site in red, CXCL10 cleavage site in yellow, GP63 binding pocket in green, and BIPSPI predicted binding sites in orange.

*In silico* modeling of the 3D protein-protein interaction using Zdock(39) demonstrated a close physical approximation of the GP63 active site catalytic zinc ion to the cleavage site on CXCL10 (Fig. 2C-F). First, GP63 and CXCL10 were loaded into Zdock with no prior information regarding binding. In the top 10 modelling predictions from this unbiased approach CXCL10 consistently localized to the binding pocket; however, no consensus emerged for the chemokine orientation within the binding pocket and the average distance from cleavage site to catalytic zinc ion was 31.92 ± 7.27 Å (Fig. 2D). Second, the Zdock modeling was repeated with the contact residues predicted by BIPSPI specified. This resulted in a consistent fit of CXCL10 into the GP63 binding pocket with the average distance between the CXCL10 cleavage site(23) and catalytic zinc ion decreased to 12.55 ± 2.52 Å (Fig. 2D). Notably, the smallest predicted distance between substrate cleavage site and the enzyme catalytic zinc ion was 8.00 Å which is consistent with the observed distance from the co-crystal structure of matrix-metalloprotease-1 and collagen (PDB: 4AUO)(40). Therefore, the predicted binding site leads to orientation of CXCL10 such that the cleavage site is accessible by the GP63 active site.

### Episodic positive selection occurred at residues surrounding the CXCL10 binding site on GP63

Given the single amino acid substitution between *Leishmania* and *Viannia* parasites (Fig. 2B) at the predicted CXCL10 binding site, we hypothesized that this region is under positive selection that potentially contributes to the phenotype of reduced CXCL10 cleavage. Previous studies have described high rates of variation and evidence of positive selection around the protease active site in the tertiary structure(32, 33). However, these studies included 6 or fewer total *Viannia* sequences and only tested for pervasive selection. In the case of *Leishmania* spp. where there is significant heterogeneity within the phylogenetic structure, this may underestimate the degree of positive selection by discounting instances where selection only occurred in a subset of the phylogeny. Therefore, using the 54 sequences identified by BlastP above, we analyzed the evolutionary pressure on *gp63* using two models: 1) the ConSurf model which generates a conservation score based on a maximum likelihood estimate of the evolutionary rate at each amino acid site(41, 42) and 2) the Mixed Effects Model of Evolution (MEME) on the HyPhy platform which tests for episodic positive selection at each amino acid residue(43). The conservation score generated by ConSurf showed extremely high conservation around known functional residues such as the active site, met-turn, and GPI-anchor (Fig. 3A). Amino acids with low conservation scores, and are therefore highly variable, correlate with the amino acids identified as under positive selection by MEME (Fig. 3A). One such peak of variation and positive selection encompasses the putative CXCL10 binding site identified here (Fig. 3A). The high variability and evidence of positive selection around the CXCL10 binding site is consistent with this region being involved in substrate binding.

**Figure 3.**
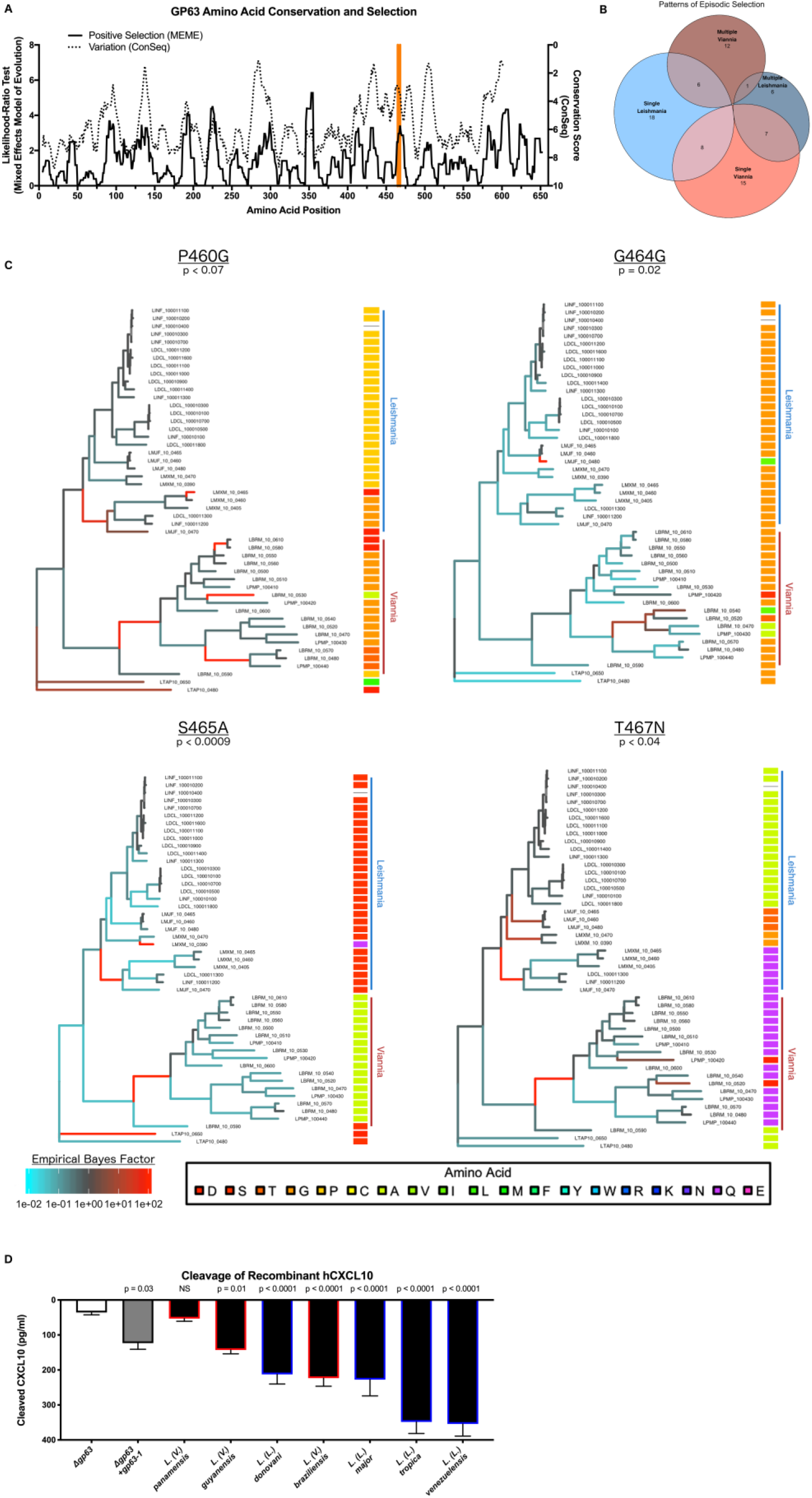
*gp63* has undergone significant positive selection between the *Leishmania* and *Viannia* subgenera, with a peak of selection identified around the CXCL10 binding site. *(A) GP63 contains multiple regions of high diversity under strong positive selection, including the region containing the CXCL10 binding site*. The conservation score was generated by the ConSeq method and ranges from 1 (highly variable) to 9 (highly conserved). Positive selection was tested at each amino acid by the likelihood ratio test from the Mixed Effects Model of Evolution (MEME). Higher likelihood ratio indicates stronger signal of positive selection. Both the conservation score and likelihood ratio test are plotted as the moving average over 10 amino acid windows. The putative CXCL10 binding site identified by BIPSPI is highlighted in orange. *(B) GP63 individual amino acids demonstrate diverse patterns of episodic selection*. The MEME test for positive selection generates an empirical bayes factor (EBF) as an estimate of the strength of positive selection for each amino acid along each branch of the phylogenetic tree. Using this measure the patterns of episodic selection were classified as single or multiple events occurring in either the *Leishmania* or *Viannia* subgenera. EBF was set to a threshold of 30 as a cutoff for positive selection. *(C) Residues adjacent to the CXCL10 binding site demonstrate a pattern of exclusive mutation between the Leishmania and Viannia subgenera*. The EBF generated by MEME was plotted onto the phylogenetic tree for positively selected residues (p<0.1) within 5 amino acids of the predicted CXCL10 binding site. The phylogenetic tree was rooted at the node of the most recent common ancestor of the two *Sauroleishmania* sequences identified. The amino acid residue for each sequence at the indicated position is plotted based on a multisequence alignment generated in ClustalOmega and used for both evolutionary tests above. *(D)* L. Viannia guyanensis *and* L. Viannia braziliensis *parasites have an intermediated CXCL10 cleavage phenotype consistent with the observation of mixed D463 and Y463 alleles in the* Viannia *subgenus*. The CXCL10 cleavage assay described in Figure 1 was repeated with *L. (L*.*) donovani, L. (L*.*) venezuelensis, L. (L*.*) tropica, L. (V*.*) guyanensis*, and *L. (V*.*) braziliensis*. 1×10^6^ parasites were incubated with 500pg/ml of human recombinant CXCL10 at 37°C for 1 hour prior to measuring the remaining CXCL10 by ELISA. Cleaved CXCL10 was calculated by subtracting the CXCL10 in the parasite conditions from a no-parasite media control. Mean ± SEM is plotted from 10 biological replicates across 5 separate experiments. P-values calculated by one-way ANOVA with Holm-Sidak test comparing each species to the *L. major Δgp63* allele.

Next, we asked whether this pattern of amino acid substitution corresponds to a difference in *L. (V*.*) panamensis* specifically or as part of a larger subset of parasites such as the *Viannia* subgenus. To do so we utilized the empirical bayes factor (EBF), a measure of the strength of positive selection generated by MEME, to examine the patterns of episodic selection within the phylogeny for each amino acid residue. The 82 amino acid sites identified as under positive selection (P < 0.05) were categorized based on whether they were positively selected along one or multiple branches within either the *Leishmania* or *Viannia* subgenera (Fig. 3B). 4 positively selected residues (p<0.1) were identified within 5 amino acid residues of the putative CXCL10 binding site. Three of these sites were mutated away from the consensus *Leishmania* subgenus residue in all sequences from the *L. (V*.*) panamensis* reference genome and in at least 17 out of 18 *Viannia* sequences analyzed: P460G, S465A, T467N (Fig. 3C). These results suggest that the change in GP63 substrate specificity is generalizable to other parasites in the *Viannia* subgenus.

### Viannia parasites demonstrate variable CXCL10 suppression corresponding to frequency of the CXCL10 specific gp63 allele

To test whether this predicted difference in substrate specificity applied to additional *Leishmania* and *Viannia* parasites, we repeated the CXCL10 cleavage assay using *L. (L*.*) donovani, L. (L*.*) venezuelensis, L. (L*.*) tropica, L. (V*.*) guyanensis*, and *L. (V*.*) braziliensis*. Similar to *L. (V) panamensis* the other two *Viannia* species tested cleaved less CXCL10 compared to the *Leishmania* species (Fig. 3D). Notably, they demonstrated an intermediate capacity to cleave CXCL10, which is consistent with the observation that unlike *L. (V*.*) panamensis, L. (V*.*) braziliensis* parasites have a mixture of *gp63* expansion copies, some of which retain the D463 allele present in the *Leishmania* subgenera while others have the *Viannia* specific N463 allele (Fig. 3C). Together this data suggests that *Viannia* parasite infection results in greater CXCL10 due to reduced GP63-mediated chemokine suppression in addition to previously described differences in host-parasite sensing.

### Mutagenesis of positively selected residues near the CXCL10 binding site on L. (L.) major gp63 to the L. (V.) panamensis sequence significantly impairs CXCL10 cleavage

We sought to experimentally test how the putative CXCL10 binding site and surrounding residues under positive selection between *Leishmania* and *Viannia* subgenera (summarized in Fig. 4A) alter the kinetics of CXCL10 cleavage by GP63. Using site directed mutagenesis of an *L. major gp63* overexpression plasmid, we mutated the CXCL10 binding site and nearest positively selected residues (Fig. 4B), expressed these constructs from HEK-293T cells, and assayed conditioned media containing equal amounts of WT and mutant GP63. *L. major gp63*^*PYSDGS*^, *gp63*^*PYS****<u>N</u>****GS*^, or *gp63*^***<u>G</u>****YS****<u>N</u>****G****<u>A</u>***^ was incubated with CXCL10 at 1000pg/ml for 2 hours to measure the cleavage rate. The wildtype *gp63*^*PYSDGS*^ has a significantly higher CXCL10 cleavage rate (6.07 pg/ml/min) than both *gp63*^*PYS****<u>N</u>****GS*^ (1.16 pg/ml/min; p=0.04) and *gp63*^***<u>G</u>****YS****<u>N</u>****G****<u>A</u>***^ (0.14pg/ml/min; p=0.01). Notably, the single mutation at position D463 predicted by BIPSPI resulted in incomplete abrogation of CXCL10 cleavage; however, in combination with mutation of the adjacent residues under positive selective pressure the cleavage rate was further reduced (Fig. 4C-D). To determine whether these mutations caused an overall loss in total proteolytic activity, all copies of *gp63* were also incubated with the non-specific substrate azocasein at 50 mg/ml and monitored over 24 hours. There was no significant difference in azocasein cleavage rate between the wildtype and either mutated copies of *gp63* (Fig. 4E-F) indicating that the observed loss of CXCL10 cleavage is not due to a reduction in total proteolytic activity. Together these data indicate that sequence variation of GP63 in the *Viannia* subgenus contributes to substrate specificity and differences in chemokine suppression. Specific changes that have been selected for in *L. panamensis* and other *Viannia* parasites markedly reduce CXCL10 cleavage while having minimal effect on overall proteolytic activity.

**Figure 4.**
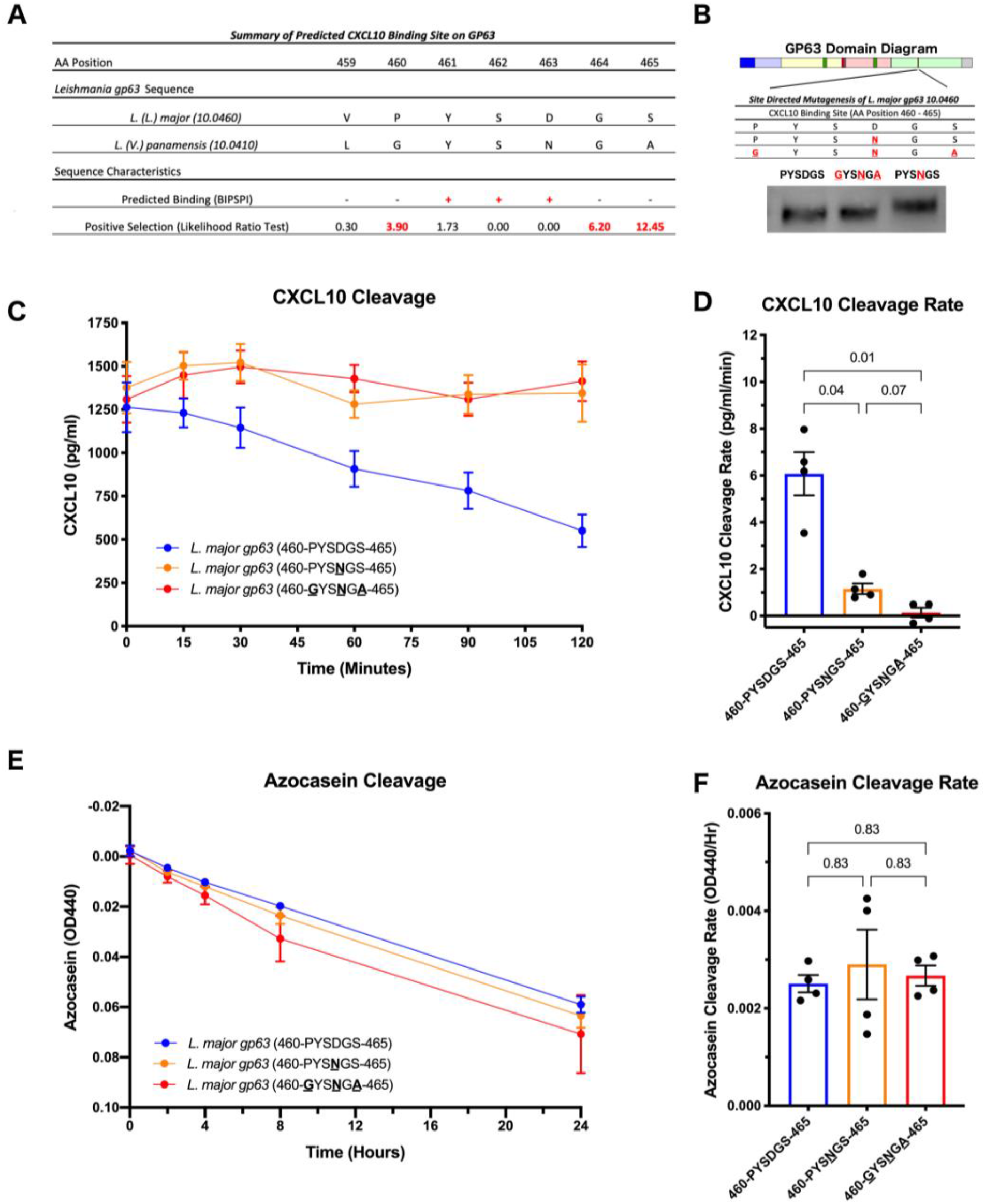
Mutagenesis of *L. (L*.*) major* CXCL10 binding site to *L. (V*.*) panamensis* residues reduces CXCL10, but not azocasein, cleavage. *(A) The predicted CXCL10 binding site is in a highly variable region of gp63 with multiple adjacent residues under positive selection*. Summary of the predicted *L. (L*.*) major* and *L. (V*.*) panamensis* CXCL10 binding alleles from BIPSPI analysis and degree of positive selection as measured by likelihood ratio test from MEME. Amino acid position number is reported relative to the L. major *gp63 10*.*0460* gene. *(B) Generation of CXCL10 binding site mutants by site-directed mutagenesis of the L. major* GP63 WT sequence. The wild-type *L. major gp63* CXCL10 binding allele (460-PYSDGS-465) cloned in an overexpression vector was mutated to the *L. panamensis* allele at the BIPSPI predicted binding site D463 (460-PYS**<u>N</u>**GS) and at the adjacent positively selected residues (**<u>G</u>**YS**<u>N</u>**G**<u>A</u>**). All three *gp63* plasmids were overexpressed in HEK293T cells and GP63 protein recovered from culture supernatants. Total GP63 protein was assessed by western blot, and samples were then diluted to equal concentration before downstream use. Representative western blot of normalized GP63 from culture supernatants is shown. (C-D) *L. (L*.*) major gp63 allele mutated to the L. (V*.*) panamensis allele at the CXCL10 binding site impairs CXCL10 cleavage*. Equal amounts of GP63 protein from each mutant were incubated with 1000pg/ml of CXCL10 for 2 hours. The remaining CXCL10 in the reaction was measured by ELISA. Change in CXCL10 over time is shown in (C) and the rate of cleavage determined by linear regression is shown in (D). (E-F) *L. (L*.*) major gp63 allele mutated to the L. (V*.*) panamensis allele at the CXCL10 binding site does not alter azocasein cleavage*. Equal amounts of GP63 protein from each mutant were incubated with 50mg/ml of azocasein for 24 hours. Cleaved azocasein was determined by measuring OD440 above background determined by the no-parasite media control Change in azocasein over time is shown in (E) and the rate of cleavage determined by linear regression is shown in (F). For (C-F) mean +/-the standard error of the mean is plotted from 4 unique experiments representing recombinant GP63 generated from two separate transfections. For D and F, P-values calculated by one-way ANOVA with holm-sidak post-hoc test.

### Within the Viannia subgenus, clinical isolates of L. (V.) braziliensis have a higher proportion of the CXCL10 cleaving GP63 D463 allele compared to L. (V.) panamensis

We next sought to determine the potential for the relative amounts of D463 and N463 alleles to contribute to variation in leishmaniasis by analyzing GP63 sequence variation from recently isolated parasites from Colombia and Bolivia (44, 45). Patino et al. sequenced whole genomes from parasites (19 *L. (V*.*) panamensis(44)* and 7 *L. (V*.*) braziliensis(45)*) from 26 infected patients. Due to the high rates of *gp63* gene duplication and copy number variation, significant genetic heterogeneity occurs among parasite isolates and structure of the region is poorly understood even in the reference PSC-1 strain. Therefore, we aligned the available sequences to the *L. (V*.*) panamensis* PSC-1 reference genome(25) and quantified the relative D463 and N463 allele frequencies based on the read depth at all mapped *gp63* positions combined. Notably, all *L. (V*.*) panamensis* isolates predominantly encode the N463 allele which does not cleave CXCL10. However, the *L. (V*.*) braziliensis* clinical isolates have greater frequency of the CXCL10 cleaving D463 allele (Fig. 5A-B). There were also additional less frequent variants that encode substitutions to alanine and threonine at this locus. The variation matched the phylogenetic analysis of *Leishmania* reference sequences (Fig. 3C) and was consistent with the intermediate CXCL10 cleavage phenotype observed with *L. (V*.*) braziliensis* (Fig. 3D). The observation that both the CXCL10 cleaving and non-cleaving allele are present in naturally occurring *Viannia* isolates confirmed the relevance of this diversity in clinical isolates and suggests that human infection likely also varies in the type and robustness of CXCL10 mediated inflammation during infection.

**Figure 5.**
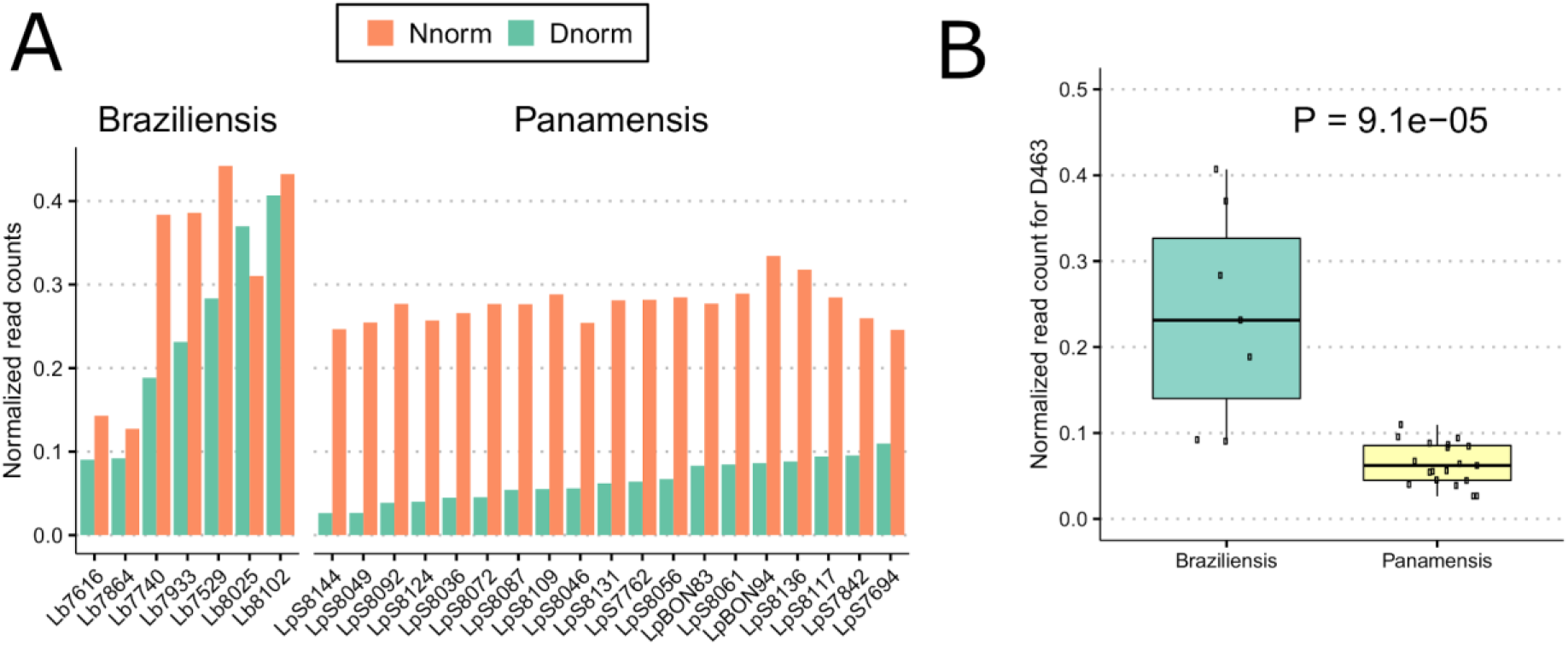
Clinical isolates of *L. (V*.*) panamensis* have predominantly CXCL10 non-cleaving D463 allele, whereas *L. (V*.*) braziliensis* have the D463 allele in addition to the CXCL10 cleaving N463. *(A) Barplot of proportion of CXCL10 cleaving (N463) and non-cleaving (D463) alleles in individual clinical isolates*. Short read sequences from 7 *L. (V*.*) braziliensis* and 19 *L. (V*.*) panamensis* were obtained from Patino et al. (2020) and Patino et al. (2020) and aligned to the *L. (V*.*) panamensis* PSC-1 reference genome. We then quantified the number of reads carrying the amino acid allele (N, D, A, or T) at *gp63* position 463. The read number was further normalized to total read depth of each sample with a scale factor of 10000, depicted as “Normalized read counts” (y axis). Amino acid position number is reported relative to the the *L. major 10*.*0460* sequence. (B) L. (V.) braziliensis *clinical isolates have significantly higher proportion of CXCL10-cleaving D463 allele than L. (V*.*) panamensis* isolates. Boxplot of the 26 clinical isolates demonstrates the distribution of D463 frequency across *L. (V*.*) braziliensis* and *L. (V*.*) panamensis*. The normalized read counts (y axis) were calculated as described in Figure 5A. P-value calculated using Wilcoxon rank-sum test.

## Discussion

Genetic diversity of *Leishmania* spp. contributes to variable disease presentation, but how the diversity alters molecular mechanisms to impact disease outcome is incompletely understood. Here we described how variation in chemokine suppression by the virulence factor *gp63* between the *Leishmania* and *Viannia* subgenera is driven by genetic variation generated by positive selection. Thus, we have identified the molecular and evolutionary basis for one difference in immune evasion between closely related parasites, and undoubtedly more remain to be discovered.

We characterized a change in substrate specificity for human CXCL10; however, the genetic variation between subgenera potentially confers loss or gain of specificity for other GP63 substrates. GP63 has a myriad of described substrates(20) with roles in host defense against infection ranging from interfering with complement mediated lysis(21) to altering intracellular signaling(46-48) to impairing antigen cross-presentation(49). Our work here shows that cross species variation significantly alters GP63 function, resulting in a change in chemokine landscape during infection. Future studies are needed to establish whether *Leishmania* diversity influences GP63 specificity and activity for other known substrates. It is also possible that mutations have led to gain of function for currently undiscovered substrates, which may alter chemokine signaling or other aspects of the immune response that contribute to the diverse clinical outcomes associated with *Leishmania* infection.

Variation in chemokine response due to GP63 diversity is likely to contribute to the severity of disease outcome by altering the balance between a well-regulated protective host immune response and dysregulated immune mediated pathology. *Leishmania major* was used as a model organism to establish the paradigm of a protective T_h_1 cell response against intracellular pathogens(5, 6). However, recent advances have described that severe, ulcerative lesions caused by *Viannia* subgenera parasites, including *L. panamensis* and *L. braziliensis*, are characterized by markedly increased expression of type-1associated cytokines and chemokines including CXCL10 (13, 50). This is known to be in part due to differences in host Toll-like receptor recognition of the parasite leading to differences in transcriptional changes(15, 26). Here we uncovered another layer of this complex host-pathogen interaction where in addition to differences in host sensing, *Viannia* parasites appear to rely on alternative mechanisms of immune evasion to their related *Leishmania* parasites. While we observed that both CXCL10 cleaving and non-cleaving alleles occurred in *L. (V*.*) panamensis* and *L. (V*.*) braziliensis* clinical isolates, the sample size and limited clinical information was insufficient to determine if the *gp63* allele was associated with specific clinical outcomes. Thus, future studies are needed to test how this diversity in parasite manipulation of chemokine signaling impacts pathogenesis in animal models and human disease. Specifically, given the significant *gp63* genetic diversity in both nucleotide and copy number variations, long-read sequencing from a large number of clinical derived samples is warranted to accurately map this region and test for associations with clinical outcomes.

The observation that genetic diversity contributes to variation in immune evasion raises the question as to what factors are responsible for generating and maintaining greater diversity in the *Viannia* subgenus. The diversification appears to be facilitated by the large copy number expansion in the *Viannia* subgenus(27, 51). It is possible that the initial gene expansion of *gp63* was driven by pressure to counteract host CXCL10 production increased in New World *Viannia* infections in part due to the presence of RNA viruses infecting either parasite or host(12-15, 17). Consistent with a model of environmental pressure in the New World driving the expansion of GP63, Bussotti et al. have described a large copy number expansion of GP63 in an *L. (L*.*) infantum* isolate from Brazil compared to isolates from the Old World(52). Such gene duplications are a common mechanism allowing for diversification of novel genes in multiple biological systems(53). By maintaining the original copy, the novel copy confers freedom to explore novel biochemical space and test new strategies for parasite survival, persistence, and spread. Eventually with a different strategy to survive in mammalian hosts, the dependence on the older mechanism of CXCL10 suppression could become expendable leading to loss of the CXCL10 specific alleles. Further, the greater diversity of *gp63* maintained within the *Viannia* subgenera may itself provide an evolutionary advantage. Within 41 *L. (V*.*) braziliensis* isolates from a single location in Brazil, 45 different polymorphic alleles were identified within *gp63*(28), and *L. (V*.*) braziliensis* isolates vary significantly in several phenotypes of immune modulation *in vitro(31)*. A larger pool of sequences conferring unique substrate specificities can allow for rapid adaptation to new environmental or host challenges. Additional studies are warranted to further characterize the evolutionary and functional implications of *gp63* diversity within the *Viannia* subgenus. Regardless of how the expansion and diversification occurred, our results clearly demonstrate that current *Leishmania* isolates have genetic variation in *gp63* that contributes to differences in host chemokine levels during infection. Further, the level of CXCL10 appears to have been selected for by the particular niche occupied by each *Leishmania* species, with variation in time and space for an optimal level of CXCL10 resulting in balancing selection and maintenance of diversity for the *Viannia* subgenus.

*Leishmania* genetic diversity creates a complex set of interactions between host and parasite, but also represents significant opportunities to leverage that diversity to improve our understanding of mechanisms of pathophysiology. Studies such as this one that link genetic variation to phenotypic differences in chemokine signaling will be required to fully understand the parasite factors that contribute to differential host susceptibility to infection. A more complete understanding of this molecular evolution will facilitate the development of biomarkers and host-directed therapies to improve outcomes of leishmaniasis.

## Materials and Methods

### Human Cell Lines and Culture

THP-1 monocytes, originally from the American Type Culture Collection (ATCC), were obtained from the Duke Cell Culture Facility and maintained in RPMI 1640 media (Invitrogen) supplemented with 10% fetal bovine serum (FBS), 2 mM glutamine, 100 U/ml penicillin-G, and 100 mg/ml streptomycin. HEK293T cells were obtained from ATCC and maintained in DMEM complete media (Invitrogen) supplemented with 10% FBS, 100 U/ml penicillin-G, and 100 mg/ml streptomycin. All cell lines were maintained at 37°C with 5% CO_2_. For phorbol 12-myristate 13-acetate (PMA) differentiation of THP-1 monocytes, 1.2 × 10^6^ cells were placed in 2 mL of complete RPMI 1640 media supplemented with 100 ng/mL of PMA for 16 hours after which the RPMI media was replaced and cells allowed to rest for 24 hours prior to infection.

### Parasite Culture and Infections

All *Leishmania* parasites were maintained in M199 supplemented with 100U/ml penicillin/streptomycin and 0.05% hemin. The following parasites strains were obtained from Biodefense and Emerging Infections (BEI) Resources or ATCC: *L. major* Δ*gp63* [(MHOM/SN/74/SD) Δ*gp63 1-7*, NR-42489], *L. major* Δ*gp63*+*1* [(MHOM/SN/74/SD) Δ*gp63 1-7* + *gp63-1*, NR-42490], *L. major* Friedlin V1 [(MHOM/IL/80/FN) NR-48815], *L. tropica* [(MHOM/AF/87/RUP) NR-48820], *L. donovani* [(MHOM/SD/62/1S) NR-48821], *L. venezuelensis* [(MHOM/VE/80/H-16) NR-29184], *L. braziliensis* [(MHOM/BR/75/M2903) ATCC-50135], *L. guyanensis* [(MHOM/BR/75M4147), ATCC-50126], and *L. panamensis* [(MHOM/PA/94/PSC-1), NR-50162]. Prior to infection parasites were washed once with HBSS and counted by hemocytometer prior to resuspending in the indicated assay media.

### Human chemokine detection

Human CXCL10 mRNA and protein were detected after infection *in vitro*. At the time of harvest, infected cells were spun at 200g for 5 minutes and the supernatants removed and stored at -80°C storage for cytokine detection. Cells were resuspended in RNAprotect Cell Reagent (Qiagen) and stored at -80°C prior to extracting RNA with RNeasy RNA extraction kit (Qiagen). Reverse transcriptase was performed using iScript Reverse Transcriptase kit (BioRad, 1708840). Quantitative real-time PCR (qRT-PCR) was performed using iTaq Universal Probes Supermix (Biorad, 1725135) with human CXCL10 (Thermo, Hs01124252) or rRNA45s5 (Thermo, Hs03928990_g1). Relative expression was calculated by the ΔΔCt method relative to the rRNA45s5 housekeeping gene. hCXCL10 protein concentration in supernatant was assayed by ELISA (R&D, 266-IP).

### Prediction and modeling of CXCL10 binding site on GP63

*In silico* prediction of the CXCL10 binding site on GP63 involved two sequential steps utilizing the amino sequence and solved crystal structures in the Protein Data Bank (GP63: 1lml and CXCL10: 1o7y). First, the primary amino acid sequences for GP63 and CXCL10 were input to the the xgBoost Interface Prediction of Specific-Partner Interactions (BIPSPI)(34) webserver (http://bipspi.cnb.csic.es/xgbPredApp/) to predict the interacting residues. Second, the crystal structures for GP63 and CXCL10 were uploaded to the Zdock webserver (http://zdock.umassmed.edu/)(39) to model the protein-protein docking interaction. The Zdock modeling was performed under two sets of conditions: 1) blinded to any knowledge of predicted contact residues and 2) with the contact residues identified by BIPSPI specified. The top ten models for the GP63-CXCL10 interaction under each conditioned were visualized using PyMol(54). Distance measurements were performed with the PyMol “distancetoatom” function relative to the catalytic zinc ion in GP63 (PDB 1lml).

### Evolutionary analysis of GP63 sequences

To examine the evolutionary pressure on GP63 we 1) identified a set of GP63 sequences to create a multisequence alignment, 2) tested for the degree of conservation at each amino acid residue, and 3) tested for episodic positive selection at individual amino acid residues. First, GP63 sequences were identified using BlastP with *gp63* from *L. major* Fd (*LmjF_10*.*0460*) used as a query (E: 0.01, no low complexity filter) on TriTrypDB(55) to search all *Leishmania* spp. The identified sequences and identifying information were downloaded. The publicly available sequences were then filtered based on the following characteristics: length (greater than half the length of the reference GP63 (302 amino acids), presence of ambiguous amino acids, genomic location (restricted to the chromosome 10 locus of GP63). The remaining 54 full length nucleotide sequences from the chromosome 10 locus were then aligned using ClustalOmega(56). Finally, using AliView(57) the alignment was manually inspected, all stop codons were removed, and the nucleotide sequence was translated to amino acid sequence. Second, the degree of conservation at each position was analyzed using the ConSurf(41) server with the ConSeq(42) method. The translated amino acid sequence created above was uploaded, the 3D structure was not specified in order to include residues from the complete peptide (the crystal structure was only solved beginning at residue 100), setting *LmjF_10*.*0460* as the query sequence, calculating the phylogenetic tree by Bayesian method. Third, episodic positive selection was tested for by the Mixed Effects Model for Evolution(43) on the Datamonkey 2.0 server(58) using the nucleotide alignment generated above. The empirical bayes factor was mapped onto the generated phylogenetic tree using the ggtree(59) package in R(60). Prior to mapping the phylogenetic tree was rooted to the node at the base of the *Sauroleishmania* subgenus using the root function in the ape package(61).

### Expression of recombinant GP63 and Site-Directed Mutagenesis

Overexpression and mutagenesis of GP63 was performed as described previously(23). In brief, 250,000 HEK293T cells were washed once with PBS, resuspended in serum free, FreeStyle 293 Expression Media (ThermoFisher, 12338018), and plated in a 6-well tissue culture treated dish 48 hours before transfection. One hour prior to transfection, media was replaced with fresh FreeStyle 293 media. Transfections were performed following the manufacturer’s protocol with the Lipofectamine 3000 transfection reagent kit. Supernatants were harvested 48 hours after transfection and stored in single use aliquots in low binding tubes at -80°C. Site directed mutagenesis was performed using the Agilent Quick Change Site Directed Mutagenesis kit per the manufacturers protocol.

### CXCL10 and azocasein cleavage assays

To assay GP63 activity two substrates were used: human CXCL10 and the non-specific colorimetric protease substrate azocasein. To normalize the total number of live parasites assayed: 8mL of a day 6 culture of promastigotes was washed once with HBSS and counted by hemocytometer to load at total of equal number of parasites for each substrate reaction. For heterologous expressed GP63, the relative amount of GP63 in each reaction was normalized based on total GP63 detected by western blot for the C-terminal histidine epitope tag. Protein was first separated by electrophoresis in a 4-20% bis-tris polyacrylamide gel before transferring to PVDF membrane using a Hoefer TE77X semi-dry transfer system. GP63 was detected by primary anti-his antibody (Cell Signaling Technology, 12698) with a secondary anti-rabbit fluorescent probe (Licor, IRDye 800CW) and developed with LiCor Odyssey Infrared Imaging System. Relative band intensity was quantified and used to produce even loading of GP63 mutants in the subsequent cleavage reaction. Dilutions of the GP63 mutants were rerun by western to confirm that equal amounts of protease were added to all reactions.

To monitor GP63 cleavage of CXCL10, normalized live parasites (1×10^6^) or heterologously expressed GP63 was incubated with 500pg/ml of human recombinant CXCL10 (Peprotech) for the indicated time at 37°C. The remaining CXCL10 was assayed by hCXCL10 enzyme-linked immunosorbent assay (ELISA) (R&D, 266-IP). The cleaved fraction of CXCL10 was calculated by subtracting the remaining measured CXCL10 from the no parasite or no transfection control. To monitor GP63 cleavage of azocasein cleavage 50 μl of normalized live parasites (5×10^7^) or heterologously expressed GP63 was incubated with 200μl of 50mg/ml of azocasein (Sigma, A2765) for the indicated time at 37°C. To stop the reaction 50μl of the reaction was added to 200μl of 5% trichloroacetic acid (TCA). The precipitate was spun at 2200g for 10 minutes, and 150μl of supernatant transferred to a clean well of a clear bottom 96-well plate and add 112.5μl of 500mM NaOH to each well. Absorbance was subsequently measured at OD440 using a BioTek microplate reader. The cleaved fraction of azocasein was calculated relative to the no parasite or no transfection control.

### Quantification of gp63 expression

To quantify *gp63* expression, mRNA was obtained from day 6 of promastigote cultures of either *L. (L*.*) major* or *L. (V*.*) panamensis*. Parasites were washed once with HBSS and counted by hemocytometer prior to resuspending at 1×10^6^ parasites in 60μl. A total of 4mL of parasites were incubated at 37°C for 1 hour to mimic the THP-1 infection conditions. At the end of 1 hour, parasites were spun at 1300g for 10 minutes and resuspended in buffer RLT from the RNeasy Mini Kit (Qiagen). Genomic DNA was removed from the sample using TurboDNase (Thermo, AM2239) per manufacturers protocol. cDNA synthesis was performed with the iScript Reverse Transcriptase kit (BioRad, 1708840). qPCR reactions were performed using the iTaq Univeral SYBR Green Mastermix (BioRad, 172-5124) with 50nm of each primer and 4μl of cDNA for gene targets or 4μl of cDNA for housekeeping genes in a final reaction volume of 10μl. Relative expression was calculated as ΔCt.

Primers for *gp63* were designed to capture all copies in the chromosome 10 locus for *L. (L*.*) major* and *L. (V*.*) panamensis*. For *L. (L*.*) major* there are four copies of *gp63* in a tandem array of which 1 copy is sufficiently divergent from the other three to require unique primers. Therefore, one set of primers were designed to amplify *LmjF_10*.*0460, LmjF_10*.*0465*, and *LmjF_10*.*0480* (Fwd: CCGTCACCCGGGCCTT, Rev: CAGCAACGAAGCATGTGCC) and a separate set of primers to amplify *LmjF_10*.*0470* (Fwd: TTGAGCGGTGGAATGAGAGG, Rev: AGTGCCATGAGAGAGAGAACT). For *L. (V*.*) panamensis* all our copies were homologous enough to utilize one set of primers for all four copies: *LPMP_100410, LPMP_100420, LPMP_100430*, and *LPMP_100440* (Fwd: CCGACTTCGTGCTGTACGTC, Rev: TGAAGCCGAGGGCGTG). Previously described primers for the α-tubulin housekeeping gene(62) were used as a control because the gene is highly conserved between *L. (L*.*) major* and *L. (V*.*) panamensis*.

### Whole genome assembly and analysis of gp63 substrate specific alleles

To quantify the natural diversity of GP63 sites involved in CXCL10 substrate specificity, we reassembled the whole genomes of 26 samples from *Viannia* subgenus parasites, including 19 *L. (V*.*) panamensis* and 7 *L. (V*.*) braziliensis* parasites. Raw data (.sra) were downloaded from two previous studies (44, 45). The module *fastq-dump* from NCBI SRA toolkit v2.10.9 (https://github.com/ncbi/sra-tools/) was used to convert the raw data to FASTQ format. BWA-MEM v0.1.17(63) was then used to align all short reads to the *L. (V*.*) panamensis* PSC-1 reference genome. The reference genome was downloaded from TriTryDB database (available at https://tritrypdb.org/tritrypdb/app/record/dataset/DS_21a844223f). On this reference genome, there are four *gp63* copies (gene names: LPMP_100410, LPMP_100420, LPMP_100430, LPMP_100440) on chromosome 10, and one distantly related *gp63*-like protein (LPMP_311850) on chromosome 31 which was not included in this study.

Sequence alignment to the reference genome was performed with minimum seed length of exact match as 19 (-k). Duplicated reads were marked and filtered by *Picard* using the *MarkDuplicates* function (*Picard* was downloaded from http://broadinstitute.github.io/picard). SAMTOOLs v1.9 (64) was then used to sort and index BAM files, and the module BCFTOOLs used to pileup, count read depth, and call variants with quality filtering score of 30 (-q=30). The final variants were stored into a VCF file. Since the alleles of a variant can have different representations, we used *vt (65)* to normalize variants, and then removed duplicate variants to avoid potential inconsistent calling bias. Next, we counted the read depth mapped to GP63 protein position 463 position. All amino acid numbering for *gp63* described here is reported relative to the *L. (L*.*) major* gene *10*.*0460*. A total of four types of amino acid codons were observed: N (AAT, AAC), D (GAT, GAC), T (ACT) and A (GCT). Then we counted the number of reads mapped to each codon for each sample. To account for differences in the sequence library size, the final values were normalized by total library size and multiplied by a scale factor of 10000.

## Acknowledgements

We are grateful to Jeffrey Bourgeois and Kyle Gibbs for thoughtful discussion and support throughout this project. ALA was supported by a Triangle Center for Evolutionary Medicine (TriCEM) graduate student fellowship and by the Burroughs Welcome Fund Graduate Diversity Enrichment Program. ATM was supported by an MGM SURE Scholarship. ALA, ATM, LW, and DCK were supported by Duke Molecular Genetics and Microbiology discretionary funds.

## References

1. Alvar J, Velez ID, Bern C, Herrero M, Desjeux P, Cano J, Jannin J, den Boer M, Team Wholc. 2012. Leishmaniasis worldwide and global estimates of its incidence. PLoS One 7:e35671.

2. Burza S, Croft SL, Boelaert M. 2018. Leishmaniasis. Lancet 392:951–970.

3. Bailey F, Mondragon-Shem K, Hotez P, Ruiz-Postigo JA, Al-Salem W, Acosta-Serrano A, Molyneux DH. 2017. A new perspective on cutaneous leishmaniasis-Implications for global prevalence and burden of disease estimates. PLoS Negl Trop Dis 11:e0005739.

4. Ponte-Sucre A, Gamarro F, Dujardin JC, Barrett MP, Lopez-Velez R, Garcia-Hernandez R, Pountain AW, Mwenechanya R, Papadopoulou B. 2017. Drug resistance and treatment failure in leishmaniasis: A 21st century challenge. PLoS Negl Trop Dis 11:e0006052.

5. Heinzel FP, Sadick MD, Holaday BJ, Coffman RL, Locksley RM. 1989. Reciprocal expression of interferon gamma or interleukin 4 during the resolution or progression of murine leishmaniasis. Evidence for expansion of distinct helper T cell subsets. J Exp Med 169:59–72.

6. Scott P, Natovitz P, Coffman RL, Pearce E, Sher A. 1988. Immunoregulation of cutaneous leishmaniasis. T cell lines that transfer protective immunity or exacerbation belong to different T helper subsets and respond to distinct parasite antigens. J Exp Med 168:1675–84.

7. Ajdary S, Alimohammadian MH, Eslami MB, Kemp K, Kharazmi A. 2000. Comparison of the immune profile of nonhealing cutaneous Leishmaniasis patients with those with active lesions and those who have recovered from infection. Infect Immun 68:1760–4.

8. Carvalho EM, Correia Filho D, Bacellar O, Almeida RP, Lessa H, Rocha H. 1995. Characterization of the immune response in subjects with self-healing cutaneous leishmaniasis. Am J Trop Med Hyg 53:273–7.

9. Castellano LR, Filho DC, Argiro L, Dessein H, Prata A, Dessein A, Rodrigues V. 2009. Th1/Th2 immune responses are associated with active cutaneous leishmaniasis and clinical cure is associated with strong interferon-gamma production. Hum Immunol 70:383–90.

10. Lee SH, Charmoy M, Romano A, Paun A, Chaves MM, Cope FO, Ralph DA, Sacks DL. 2018. Mannose receptor high, M2 dermal macrophages mediate nonhealing Leishmania major infection in a Th1 immune environment. J Exp Med 215:357–375.

11. Anderson CF, Mendez S, Sacks DL. 2005. Nonhealing infection despite Th1 polarization produced by a strain of Leishmania major in C57BL/6 mice. J Immunol 174:2934–41.

12. Novais FO, Carvalho LP, Graff JW, Beiting DP, Ruthel G, Roos DS, Betts MR, Goldschmidt MH, Wilson ME, de Oliveira CI, Scott P. 2013. Cytotoxic T cells mediate pathology and metastasis in cutaneous leishmaniasis. PLoS Pathog 9:e1003504.

13. Novais FO, Carvalho LP, Passos S, Roos DS, Carvalho EM, Scott P, Beiting DP. 2015. Genomic profiling of human Leishmania braziliensis lesions identifies transcriptional modules associated with cutaneous immunopathology. J Invest Dermatol 135:94–101.

14. Bacellar O, Lessa H, Schriefer A, Machado P, Ribeiro de Jesus A, Dutra WO, Gollob KJ, Carvalho EM. 2002. Up-regulation of Th1-type responses in mucosal leishmaniasis patients. Infect Immun 70:6734–40.

15. Ives A, Ronet C, Prevel F, Ruzzante G, Fuertes-Marraco S, Schutz F, Zangger H, Revaz-Breton M, Lye LF, Hickerson SM, Beverley SM, Acha-Orbea H, Launois P, Fasel N, Masina S. 2011. Leishmania RNA virus controls the severity of mucocutaneous leishmaniasis. Science 331:775–8.

16. Rossi M, Castiglioni P, Hartley MA, Eren RO, Prevel F, Desponds C, Utzschneider DT, Zehn D, Cusi MG, Kuhlmann FM, Beverley SM, Ronet C, Fasel N. 2017. Type I interferons induced by endogenous or exogenous viral infections promote metastasis and relapse of leishmaniasis. Proc Natl Acad Sci U S A 114:4987–4992.

17. Crosby EJ, Clark M, Novais FO, Wherry EJ, Scott P. 2015. Lymphocytic Choriomeningitis Virus Expands a Population of NKG2D+CD8+ T Cells That Exacerbates Disease in Mice Coinfected with Leishmania major. J Immunol 195:3301–10.

18. Kaye P, Scott P. 2011. Leishmaniasis: complexity at the host-pathogen interface. Nat Rev Microbiol 9:604–15.

19. Gupta G, Oghumu S, Satoskar AR. 2013. Mechanisms of immune evasion in leishmaniasis. Adv Appl Microbiol 82:155–84.

20. Olivier M, Atayde VD, Isnard A, Hassani K, Shio MT. 2012. Leishmania virulence factors: focus on the metalloprotease GP63. Microbes Infect 14:1377–89.

21. Joshi PB, Kelly BL, Kamhawi S, Sacks DL, McMaster WR. 2002. Targeted gene deletion in Leishmania major identifies leishmanolysin (GP63) as a virulence factor. Mol Biochem Parasitol 120:33–40.

22. Corradin S, Ransijn A, Corradin G, Roggero MA, Schmitz AA, Schneider P, Mauel J, Vergeres G. 1999. MARCKS-related protein (MRP) is a substrate for the Leishmania major surface protease leishmanolysin (gp63). J Biol Chem 274:25411–8.

23. Antonia AL, Gibbs KD, Trahair ED, Pittman KJ, Martin AT, Schott BH, Smith JS, Rajagopal S, Thompson JW, Reinhardt RL, Ko DC. 2019. Pathogen Evasion of Chemokine Response Through Suppression of CXCL10. Front Cell Infect Microbiol 9:280.

24. Peacock CS, Seeger K, Harris D, Murphy L, Ruiz JC, Quail MA, Peters N, Adlem E, Tivey A, Aslett M, Kerhornou A, Ivens A, Fraser A, Rajandream MA, Carver T, Norbertczak H, Chillingworth T, Hance Z, Jagels K, Moule S, Ormond D, Rutter S, Squares R, Whitehead S, Rabbinowitsch E, Arrowsmith C, White B, Thurston S, Bringaud F, Baldauf SL, Faulconbridge A, Jeffares D, Depledge DP, Oyola SO, Hilley JD, Brito LO, Tosi LR, Barrell B, Cruz AK, Mottram JC, Smith DF, Berriman M. 2007. Comparative genomic analysis of three Leishmania species that cause diverse human disease. Nat Genet 39:839–47.

25. Llanes A, Restrepo CM, Del Vecchio G, Anguizola FJ, Lleonart R. 2015. The genome of Leishmania panamensis: insights into genomics of the L. (Viannia) subgenus. Sci Rep 5:8550.

26. Ibraim IC, de Assis RR, Pessoa NL, Campos MA, Melo MN, Turco SJ, Soares RP. 2013. Two biochemically distinct lipophosphoglycans from Leishmania braziliensis and Leishmania infantum trigger different innate immune responses in murine macrophages. Parasit Vectors 6:54.

27. Valdivia HO, Scholte LL, Oliveira G, Gabaldon T, Bartholomeu DC. 2015. The Leishmania metaphylome: a comprehensive survey of Leishmania protein phylogenetic relationships. BMC Genomics 16:887.

28. Medina LS, Souza BA, Queiroz A, Guimaraes LH, Lima Machado PR, E MC, Wilson ME, Schriefer A. 2016. The gp63 Gene Cluster Is Highly Polymorphic in Natural Leishmania (Viannia) braziliensis Populations, but Functional Sites Are Conserved. PLoS One 11:e0163284.

29. Rogers MB, Hilley JD, Dickens NJ, Wilkes J, Bates PA, Depledge DP, Harris D, Her Y, Herzyk P, Imamura H, Otto TD, Sanders M, Seeger K, Dujardin JC, Berriman M, Smith DF, Hertz-Fowler C, Mottram JC. 2011. Chromosome and gene copy number variation allow major structural change between species and strains of Leishmania. Genome Res 21:2129–42.

30. Victoir K, Arevalo J, De Doncker S, Barker DC, Laurent T, Godfroid E, Bollen A, Le Ray D, Dujardin JC. 2005. Complexity of the major surface protease (msp) gene organization in Leishmania (Viannia) braziliensis: evolutionary and functional implications. Parasitology 131:207–14.

31. da Silva Vieira T, Arango Duque G, Ory K, Gontijo CM, Soares RP, Descoteaux A. 2019. Leishmania braziliensis: Strain-Specific Modulation of Phagosome Maturation. Front Cell Infect Microbiol 9:319.

32. Ma L, Chen K, Meng Q, Liu Q, Tang P, Hu S, Yu J. 2011. An evolutionary analysis of trypanosomatid GP63 proteases. Parasitol Res 109:1075–84.

33. Alvarez-Valin F, Tort JF, Bernardi G. 2000. Nonrandom spatial distribution of synonymous substitutions in the GP63 gene from Leishmania. Genetics 155:1683–92.

34. Sanchez-Garcia R, Sorzano COS, Carazo JM, Segura J. 2019. BIPSPI: a method for the prediction of partner-specific protein-protein interfaces. Bioinformatics 35:470–477.

35. Cerda-Costa N, Gomis-Ruth FX. 2014. Architecture and function of metallopeptidase catalytic domains. Protein Sci 23:123–44.

36. McGwire BS, Chang KP. 1996. Posttranslational regulation of a Leishmania HEXXH metalloprotease (gp63). The effects of site-specific mutagenesis of catalytic, zinc binding, N-glycosylation, and glycosyl phosphatidylinositol addition sites on N-terminal end cleavage, intracellular stability, and extracellular exit. J Biol Chem 271:7903–9.

37. Macdonald MH, Morrison CJ, McMaster WR. 1995. Analysis of the active site and activation mechanism of the Leishmania surface metalloproteinase GP63. Biochim Biophys Acta 1253:199–207.

38. Button LL, Wilson G, Astell CR, McMaster WR. 1993. Recombinant Leishmania surface glycoprotein GP63 is secreted in the baculovirus expression system as a latent metalloproteinase. Gene 134:75–81.

39. Pierce BG, Wiehe K, Hwang H, Kim BH, Vreven T, Weng Z. 2014. ZDOCK server: interactive docking prediction of protein-protein complexes and symmetric multimers. Bioinformatics 30:1771–3.

40. Nash A, Birch HL, de Leeuw NH. 2017. Mapping intermolecular interactions and active site conformations: from human MMP-1 crystal structure to molecular dynamics free energy calculations. J Biomol Struct Dyn 35:564–573.

41. Ashkenazy H, Abadi S, Martz E, Chay O, Mayrose I, Pupko T, Ben-Tal N. 2016. ConSurf 2016: an improved methodology to estimate and visualize evolutionary conservation in macromolecules. Nucleic Acids Res 44:W344–50.

42. Berezin C, Glaser F, Rosenberg J, Paz I, Pupko T, Fariselli P, Casadio R, Ben-Tal N. 2004. ConSeq: the identification of functionally and structurally important residues in protein sequences. Bioinformatics 20:1322–4.

43. Murrell B, Wertheim JO, Moola S, Weighill T, Scheffler K, Kosakovsky Pond SL. 2012. Detecting individual sites subject to episodic diversifying selection. PLoS Genet 8:e1002764.

44. Patino LH, Munoz M, Muskus C, Mendez C, Ramirez JD. 2020. Intraspecific Genomic Divergence and Minor Structural Variations in Leishmania (Viannia) panamensis. Genes (Basel) 11.

45. Patino LH, Munoz M, Cruz-Saavedra L, Muskus C, Ramirez JD. 2020. Genomic Diversification, Structural Plasticity, and Hybridization in Leishmania (Viannia) braziliensis. Front Cell Infect Microbiol 10:582192.

46. Halle M, Gomez MA, Stuible M, Shimizu H, McMaster WR, Olivier M, Tremblay ML. 2009. The Leishmania surface protease GP63 cleaves multiple intracellular proteins and actively participates in p38 mitogen-activated protein kinase inactivation. J Biol Chem 284:6893–908.

47. Gregory DJ, Godbout M, Contreras I, Forget G, Olivier M. 2008. A novel form of NF-kappaB is induced by Leishmania infection: involvement in macrophage gene expression. Eur J Immunol 38:1071–81.

48. Gomez MA, Contreras I, Halle M, Tremblay ML, McMaster RW, Olivier M. 2009. Leishmania GP63 alters host signaling through cleavage-activated protein tyrosine phosphatases. Sci Signal 2:ra58.

49. Matheoud D, Moradin N, Bellemare-Pelletier A, Shio MT, Hong WJ, Olivier M, Gagnon E, Desjardins M, Descoteaux A. 2013. Leishmania evades host immunity by inhibiting antigen cross-presentation through direct cleavage of the SNARE VAMP8. Cell Host Microbe 14:15–25.

50. Ramirez C, Diaz-Toro Y, Tellez J, Castilho TM, Rojas R, Ettinger NA, Tikhonova I, Alexander ND, Valderrama L, Hager J, Wilson ME, Lin A, Zhao H, Saravia NG, McMahon-Pratt D. 2012. Human macrophage response to L. (Viannia) panamensis: microarray evidence for an early inflammatory response. PLoS Negl Trop Dis 6:e1866.

51. Urrea DA, Duitama J, Imamura H, Alzate JF, Gil J, Munoz N, Villa JA, Dujardin JC, Ramirez-Pineda JR, Triana-Chavez O. 2018. Genomic Analysis of Colombian Leishmania panamensis strains with different level of virulence. Sci Rep 8:17336.

52. Bussotti G, Gouzelou E, Cortes Boite M, Kherachi I, Harrat Z, Eddaikra N, Mottram JC, Antoniou M, Christodoulou V, Bali A, Guerfali FZ, Laouini D, Mukhtar M, Dumetz F, Dujardin JC, Smirlis D, Lechat P, Pescher P, El Hamouchi A, Lemrani M, Chicharro C, Llanes-Acevedo IP, Botana L, Cruz I, Moreno J, Jeddi F, Aoun K, Bouratbine A, Cupolillo E, Spath GF. 2018. Leishmania Genome Dynamics during Environmental Adaptation Reveal Strain-Specific Differences in Gene Copy Number Variation, Karyotype Instability, and Telomeric Amplification. mBio 9.

53. Taylor JS, Raes J. 2004. Duplication and divergence: the evolution of new genes and old ideas. Annu Rev Genet 38:615–43.

54. The PyMOL Molecular Graphics System VS, LLC.

55. Aslett M, Aurrecoechea C, Berriman M, Brestelli J, Brunk BP, Carrington M, Depledge DP, Fischer S, Gajria B, Gao X, Gardner MJ, Gingle A, Grant G, Harb OS, Heiges M, Hertz-Fowler C, Houston R, Innamorato F, Iodice J, Kissinger JC, Kraemer E, Li W, Logan FJ, Miller JA, Mitra S, Myler PJ, Nayak V, Pennington C, Phan I, Pinney DF, Ramasamy G, Rogers MB, Roos DS, Ross C, Sivam D, Smith DF, Srinivasamoorthy G, Stoeckert CJ Jr.,, Subramanian S, Thibodeau R, Tivey A, Treatman C, Velarde G, Wang H. 2010. TriTrypDB: a functional genomic resource for the Trypanosomatidae. Nucleic Acids Res 38:D457–62.

56. McWilliam H, Li WZ, Uludag M, Squizzato S, Park YM, Buso N, Cowley AP, Lopez R. 2013. Analysis Tool Web Services from the EMBL-EBI. Nucleic Acids Research 41:W597–W600.

57. Larsson A. 2014. AliView: a fast and lightweight alignment viewer and editor for large datasets. Bioinformatics 30:3276–8.

58. Weaver S, Shank SD, Spielman SJ, Li M, Muse SV, Kosakovsky Pond SL. 2018. Datamonkey 2.0: A Modern Web Application for Characterizing Selective and Other Evolutionary Processes. Mol Biol Evol 35:773–777.

59. Guangchuang Yu DS, Huachen Zhu, Yi Guan, Tommy Tsan-Yuk Lam. 2017. ggtree: an R package for visualization and annotation of phylogenetic trees with their covariates and other associated data. Methods in Ecology and Evolution.

60. Anonymous. R Core Team (2019). R: A language and environment for statistical computing. R Foundation for Statistical Computing, Vienna, Austria. https://www.R-project.org/.

61. Paradis E, Claude J, Strimmer K. 2004. APE: Analyses of Phylogenetics and Evolution in R language. Bioinformatics 20:289–90.

62. Pereira BA, Britto C, Alves CR. 2012. Expression of infection-related genes in parasites and host during murine experimental infection with Leishmania (Leishmania) amazonensis. Microb Pathog 52:101–8.

63. Li H. 2013. Aligning sequence reads, clone sequences and assembly contigs with BWA-MEM. arXiv:1303.3997.

64. Li H, Handsaker B, Wysoker A, Fennell T, Ruan J, Homer N, Marth G, Abecasis G, Durbin R, Genome Project Data Processing S. 2009. The Sequence Alignment/Map format and SAMtools. Bioinformatics 25:2078–9.

65. Tan A, Abecasis GR, Kang HM. 2015. Unified representation of genetic variants. Bioinformatics 31:2202–4.

